# Engineering CD3 Subunits with Endoplasmic Reticulum Retention Signal Facilitates Allogeneic CAR T Cell Production

**DOI:** 10.1101/2024.09.22.614328

**Authors:** Hamidreza Ebrahimiyan, Ali Sayadmanesh, Mahdi Hesaraki, Marzieh Ebrahimi, Hossein Baharand, Mohsen Basiri

## Abstract

The success of autologous CAR T cell therapies has driven interest in developing off-the-shelf allogeneic CAR T cells as a scalable and readily available option for broader patient access. Most of the current approaches involve the knockout of T cell receptor (TCR) subunits via genome editing for preventing graft-versus-host disease (GvHD). However, clinical translation of these methods faces challenges due to manufacturing complexities and emerging safety concerns like unintended long deletions and chromosomal loss. In this study, we explored an alternative approach by engineering synthetic CD3 subunits containing an endoplasmic reticulum retention (ERR) signal to suppress TCR surface expression by disrupting its trafficking to the plasma membrane. We screened multiple CD3-ERR candidate designs to identify the construct with the highest efficacy in TCR downregulation. The selected candidate, CD3ζ-ERR, was further characterized, demonstrating its ability to minimize TCR-mediated activation and alloreactivity without affecting T cell phenotype, cell cycle and cytokine-induced expansion. Subsequent assays revealed that CD3ζ-ERR CD19 CAR T cells retained their CAR-mediated cytotoxic function against CD19^+^ malignant cells. This study presents an alternative approach for TCR downregulation that circumvents genome editing. By using a transgene compatible with conventional viral vector delivery, this approach holds promise for scalable clinical-grade manufacturing of allogeneic CAR T cell therapies.

**Translational Impact Statement:** Our study introduces a scalable method to engineer allogeneic CAR T cells by reducing TCR expression without genome editing, thereby minimizing the risk of immune rejection (GvHD) while maintaining antitumor efficacy. This approach offers a practical and clinically translatable solution for producing “off-the-shelf” CAR T cell therapies, potentially broadening access to these life-saving treatments and streamlining their integration into existing clinical manufacturing processes.

## Introduction

Chimeric antigen receptor (CAR) T cell immunotherapy has emerged as a groundbreaking approach in the treatment of advanced cancers, especially hematologic malignancies. The clinical success of CAR T cell therapies has led to the approval of several autologous CAR T cell products, wherein a patient’s own T cells are genetically engineered to express a CAR that specifically targets cancer antigens.^1, 2^ While these autologous CAR T cell therapies have shown remarkable efficacy, their broad clinical application is limited by several critical challenges. These include high production costs, potential contamination of the therapeutic cell product with malignant cells, variability in product quality, and a significant risk of manufacturing failure, especially in heavily pre-treated patients.^3-5^ Additionally, the time-intensive nature of manufacturing autologous CAR T cells can lead to treatment delays, which can be problematic in aggressive cancers.^6^ To address these limitations, there is growing interest in the development of allogeneic CAR T cell products, which are derived from healthy donors and can be produced in large quantities as “off-the-shelf” products. Theoretically, allogeneic CAR T cells offer several advantages, including reduced production costs, increased product uniformity, standardization of manufacturing processes, and the elimination of delays associated with the need to generate patient-specific therapies.

However, the development of allogeneic CAR T cell therapies faces its own set of challenges. One of the most significant challenges in developing allogeneic CAR T cell therapies is the risk of Graft-versus-Host Disease (GvHD), which arises from the immunologic mismatch between donor CAR T cells and the recipient.^3^ Gene editing is a commonly employed strategy to mitigate this risk by disrupting the endogenous expression of T cell receptor (TCR) subunits, such as TCR-α and CD3ε.^7^ Genome editing technologies, including CRISPR-Cas9, have proven effective in targeting multiple genes within T cells, and several genome-edited adoptive T cell therapies, including allogeneic TCR-knockout CAR T cells, are currently undergoing clinical trials.^8^ However, the large-scale production of clinical-grade genome-edited CAR T cells is technically complex and requires extensive quality control to ensure the precision of the genetic modifications.^9^ Additionally, recent studies have raised concerns about unintended large deletion^10^, chromosomal inversion^11^, and partial or complete chromosome loss^12^ following genome editing in T cells, underscoring the need for more stringent manufacturing and safety assessments.

Given these challenges, alternative approaches to downregulate TCR expression without the use of genome editing technologies are gaining attention. One such approach involves the use of RNA interference to suppress endogenous TCR expression in T cells.^13^ Another strategy employs a chimeric protein expression blocker (PEBL), which utilizes an anti-CD3 single-chain variable fragment to sequester the TCR complex, thereby preventing its expression on the cell surface.^14^ However, further research is required to refine these strategies and expand additional engineering solutions to produce safe and effective allogeneic CAR T cells.

In the present study, we sought to expand the toolkit for allogeneic CAR T cell development by devising an alternative genome-editing-free method to downregulate TCR surface expression in T cells. Our design involves the addition of an endoplasmic reticulum retention (ERR) signal to the C-terminus of CD3 subunits to interfere with the transport of the TCR complex to the cell membrane, thereby reducing its surface expression. We investigated the impact of these synthetic CD3-ERR transgenes on TCR expression and evaluated its potential for facilitating the generation of allogeneic CAR T cells.

## Methods and materials

### Cell lines and culture conditions

Platinum-A (Plat-A) and PG13 (ATCC CRL-10686) cell lines were cultured in IMDM (Iscove’s Modified Dulbecco’s Medium, Gibco cat# 21980032) with 10% HyClone Characterized Fetal Bovine Serum (FBS) (Cytiva cat# SH30071.03IH30-45) and 2 mM GlutaMax (100x, Gibco cat# 35050-061) that incubated at 37°C in 5% CO2. Firefly luciferase (ffluc)-expressing Raji cells (Royan Institute Cell Bank) were cultured in RPMI-1640 (Cytvia cat# SH3009601) with 10% HyClone FBS and 2 mM GlutaMax.

### Vector constructs and retrovirus production

The DNA sequence coding CD3ζ and CD3ε with ERR was synthesized and received in pUC57 plasmid. The tCD3ζ-ERR and tCD3ε-ERR (truncated CD3) sequences were generated by eliminating the intracellular domain of CD3 using SOEing PCR and cloned into SFG-IRES-eGFP retroviral plasmid using NcoI and SphI restriction sites. Coding sequences were confirmed by Sanger sequencing.

Plat-A cells were transiently transfected with CD3ζ-ERR, CD3ε-ERR, tCD3ζ-ERR, tCD3ε-ERR and control plasmids by Lipofectamine 3000 (Invitrogen, Part no: 100022234) according to the manufacturer’s protocol. The supernatants, containing retroviral particles, were collected and filtered (0.45 µm) after 24 hours. These viruses were used to transduce PG13 cells to generate permanent retrovirus producer cell lines. Plat-A virus supernatants supplemented with 10 μg/mL polybrene (MERK, SKU: TR-1003-G) were used for PG13 transduction. The transduction procedure was repeated until 80% of PG13 were GFP^+^ (GFP positive). Then GFP^+^ cells were flow sorted to be used as a producer cell line. The PG13 producer cell lines were seeded (43 × 10^3^ cell/cm^2^) and the media was replaced after 48 hours. 24 hours later, retroviral supernatants were collected, filtered and snap-frozen on dry ice. The vials containing the viruses were stored at −80°C for T cell transduction. The CD19 CAR virus was sourced from the Royan Institute virus lab.^15^

### T cell transduction and expansion

Peripheral blood samples were obtained from healthy donors after obtaining informed consent according to protocols approved by the Ethics Committee of the Royan Institute. Peripheral blood mononuclear cells (PBMCs) were isolated by centrifugation using Lymphoprep (Serumwerk Bernburg, cat# 1114544) and activated on anti-CD3 and anti-CD28 (Miltenyibiotec, cat# 130-093-387/ 130-093-375) pre-coated 24 well not-treated plates in CTL media containing 45% RPMI-1640, 45% Click’s medium (Irvine Scientific cat# 9195), 10% FBS and 2 mM GlutaMAX). The next day Interleukin-2 (IL-2; R&D cat# 202-IL-010/CF) was added to the final concentration of 100 U/mL, followed by 100 U/mL every other day to maintain the culture. On day 5, T cells were washed, and the medium was replaced with Human Platelet Lysates (HPL) 2.5% (Cell Tech Inc, Iran). On days 2 and 3, 2.5 ml of retroviral supernatant was transferred to a non-treated 24-well plate pre-coated with RetroNectin reagent (Takara cat# T100A/B) and centrifuged at 2000 g at 4°C for 2 hours to transduce T cells. After centrifugation, the supernatants were removed from the 24-well plate. Activated T cells (2×10^5^ cells) were resuspended in CTL media containing IL-2 (100 IU/mL), transferred to the plate, and centrifuged at 400 g for 10 min.^16^

### Flow cytometry analysis and cell sorting

For staining the cell surface markers, 2×10^5^ cells were washed and stained with specific antibodies (Table 1) at 4°C for 45 minutes in the dark. For intracellular staining of CD3 or Ki67, the fixation procedure was conducted according to the manufacturer’s protocol (BD Perm/Wash cat# 51-2091KZ) and stained at 4°C for 60 min in the dark. For cell cycle analysis,^17^ 1×10^6^ cells were fixed and permeabilized with 70% ethanol at −20°C in a drop-wise manner and incubated overnight at −20°C. The cells were washed two times with FACS buffer (1X PBS, 2% v/v heat-inactivated FBS, 1 mM EDTA), followed by PI staining (1X PBS, 100 μg/ml RNase, 50 μg/ml PI, 2 mM MgCl2) at 25°C for 20 min in the dark. All samples were acquired on BD FACSCalibur or BD FACSAria II flow cytometers (BD Biosciences, Franklin Lakes, NJ, USA). Analysis was performed using FlowJo software (Tree Star Inc., Ashland, USA).

**Table 1.** Antibodies for flow cytometry.

| Target antigen | Fluorophore | Company | Cat. No. |
| --- | --- | --- | --- |
| CD3 | PerCP | BioLegend | 344813 |
| CD8 | PerCP | BioLegend | 344707 |
| CD69 | PE | BioLegend | 310905 |
| Ki67 | PerCP-Cy5.5 | BioLegend | 350520 |

To evaluate suppression of cell surface CD3 expression while correcting for variations in transgene expression levels, and CD3 staining intensity across different groups, a normalized suppression index (NSI) was calculated using geometric mean fluorescent intensity MFI of GFP and CD3 staining in transduced (GFP^+^) and untransduced (GFP^−^) subpopulation in each sample by the following formula:

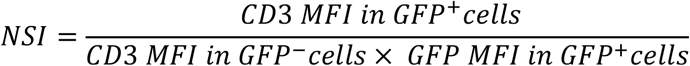

For fluorescence-activated cell sorting (FACS), 2×10^6^ cells were stained as described above and sorted using a BD FACSAria II flow cytometer (BD Biosciences, Franklin Lakes, NJ, USA). In the control group, GFP^+^ cells were gated for sorting. However, for either CD3ζ-ERR T cells or CAR CD3ζ-ERR T cells, FACS sorting was done using GFP^+^ CD3^−^.

### Assessment of proliferation and survival of CD3ζ-ERR T cells

To analysis CD3ζ-ERR toxicity in T cells, 2×10^5^ cells were seeded in two wells of a 24-well plate on day 7. On day 9, T cells were counted and seeded again as same as on day 7. T cells counting, cell cycle analysis and CD8 labeling were done on day 11. Cell viability was also assessed using PI labeling.

### Assessment of T cell activation

Sorted transduced T cells (GFP^+^CD3^−^ CD3ζ-ERR T cells and GFP^+^ control T cells) were rested in CTL medium without IL-2 for 12 hours. 2 ×10^5^ sorted T cells were then seeded in a 96-well plate pre-coated with agonistic anti-CD3 (clone OKT3, Miltenyibiotec). After 48 hours, T cells were collected and the expression of CD69 and CD3 were measured by flow cytometry. The supernatants were preserved at −20°C until they were assessed for IL-2 concentration using Human IL-2 DuoSet ELISA kit (DY202, R&D Systems) according to the manufacturer’s instructions.

### In vitro alloreactivity of CD3ζ-ERR T cell

PBMCs (10^6^ cells) obtained from healthy donors were treated with mitomycin C (20 µg/ml) to arrest the proliferation in mitomycin C at 37°C for 1 hour in the dark. Sorted T cells (2×10^5^) rested without IL-2 for 12 hours were cultured with mitomycin C-treated polled allogeneic PBMCs (4 × 10^5^) in a 24-well plate. After 72 hours of co-culture, intracellular staining for Ki67 was used to measure proliferation T cells. The supernatants were stored at −20°C until they were assessed for IFN-γ secretion. The Human IFN-γ DuoSet ELISA kit (DY285B, R&D Systems) was used to measure IFN-γ according to the manufacturer’s instructions.

### CARCD3ζ-ERR T cell generation and cytotoxicity assay

Sorted transduced T cells were rested in CTL without IL-2 for 12 hours. In a 96-well plate, 1×10^5^ T Cells were co-cultured with Firefly luciferase-expressing Raji cells in effector to target ratio (E:T) ratios of 1:1, 10:1, and 20:1 for 24 hours subjected to flow cytometry assessment of CD69 activation marker expression. In order to measure cytotoxicity, D-Luciferin (IVISbrite D-Luciferin Potassium Salt, Revvity Part Number: 122799) was used to measure the luminescence signal in each well according to manufactures protocol. The luminescence signal in each well was measured using UVTEC Cambridge. The supernatants were kept at −20°C and assessed for IFN-γ secretion using the Human IFN-γ DuoSet ELISA kit (DY285B, R&D Systems) according to the manufacturer’s instructions.

### Statistical analyses

The GraphPad Prism version 9 software (GraphPad Software, La Jolla, CA) was used for plotting and calculating data. One-way ANOVA or unpaired t test were applied based on groups. The mean ± STD was expressed as scale data, and P < 0.05 was defined as a statistically significant level. In the cytotoxicity assay, the positive results were quantified with ImageJ software V18. The quantified results were normalized based on the average of the non-transduced T cell signal, which was applied as 100.

## Results

### Screening CD3-ERR candidates for transduction efficiency and CD3 suppression

The TCR complex is transported to the cell surface only after the complete assembly of all subunits in the endoplasmic reticulum (ER). Leveraging this principle, we engineered ERR signal containing CD3 subunits, which integrate into the forming endogenous TCR complexes in the ER, thereby preventing their translocation to the cell surface (Figure 1A). We focused on CD3ε and CD3ζ because of their higher stoichiometry (two copies each) compared to other TCR subunits (one copy each), suggesting a greater likelihood of incorporating transgenic modified subunits into endogenous TCR complexes. In our four candidate retroviral constructs, ERR sequence was fused to the c-terminal cytoplasmic part of either full-length (CD3ζ-ERR and CD3ε-ERR) or truncated forms (tCD3ζ-ERR and tCD3ε-ERR) of these subunits (Figure 1B). Five days after transduction of T cells with the constructs (Figure 1C), all four candidates showed different levels of surface CD3 downregulation, which inversely correlated with GFP intensity (Figure 1D). The frequency of the CD3^−^ cells within the transduced (GFP^+^) population was significantly higher in CD3ζ-ERR T cells compared to GFP-FFluc-transduced control group (Ctrl) and all other CD3-ERR T cells (Figure 1E). Notably, CD3ζ-ERR T cells transduction rate (GFP^+^) was also the highest among the CD3-ERR groups and comparable with the Ctrl group while other candidates, especially tCD3ε-ERR, showed relatively lower transduction efficiencies (Figure 1F). Since CD3 downregulation correlated with transgene expression, as indicated by GFP signal intensity, we normalized the CD3 reduction (relative to the CD3 signal in the untransduced subpopulation) by GFP intensity. This normalization allowed us to compare the intrinsic suppression capacity of the CD3-ERR candidates, independent of variations in transduction efficiency, which was found to be comparable across all four candidates (Figure 1G). Given that CD3ζ-ERR consistently showed the highest transduction efficiency, resulting in the largest CD3^−^ population, this construct was selected for further investigation throughout the study.

**Figure 1.**
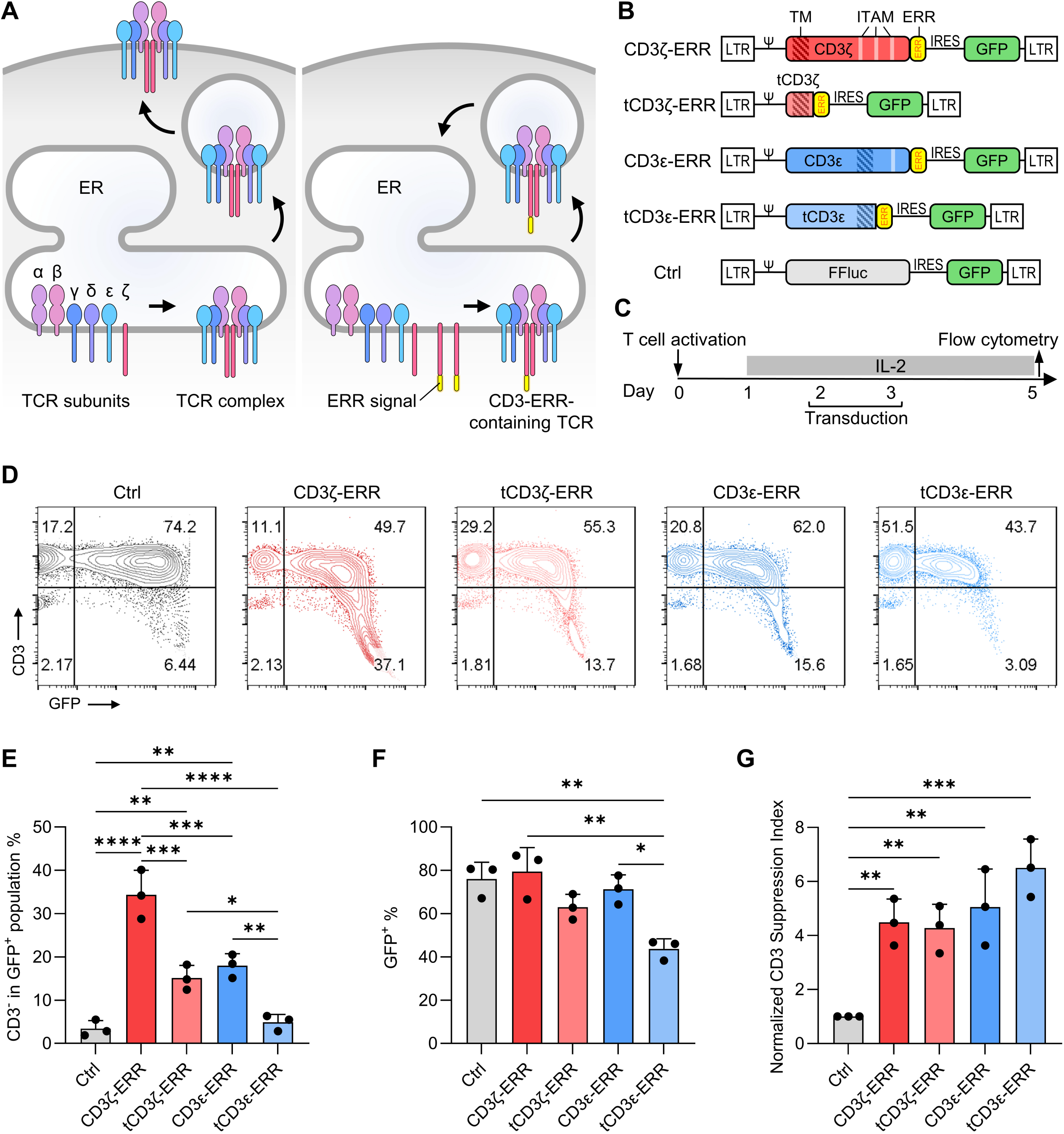
Engineered T cells, vector selection. **A**.CD3-ERR competes with endogenous CD3 in TCR complex incorporation, which leads to the blocking transportation in the case of CD3-ERR TCR complex. **B**. Schema of CD3ζ-ERR, tCD3ζ-ERR, CD3ε-ERR, tCD3ε-ERR and Ctrl. All vectors have ERR signal peptide following IRES GFP. **C**. The schematic timeline of the experimental setting. PBMC was activated by anti-CD3/CD28 on day 0. The all constructs were transduced into T cells on day 2 and 3. Flow analysis applied on day 5. **D**. The scatter plot represents the GFP and CD3 expression in different vectors. **E**. The bar graph represents the percentages of GFP^+^ T cell populations on day 5, CD3ζ-ERR have no significant difference with Ctrl. **F**. The bar graph represents the percentages of GFP^+^ CD3^-^ T cell populations on day 5, CD3ζ-ERR have significant difference with all other vectors and also Ctrl. **G**. The normalized CD3 suppression index ratio revealed no significant difference between different vectors. The data are presented as mean ± SD, \*\*\*\**P* < 0.0001, *** *P* < 0.001, ** *P* < 0.01, * *P* < 0.05, one-way ANOVA was applied for statistical analysis.

### CD3ζ-ERR suppress TCR expression on T cell surface by containing it in the intracellularly

To investigate the mechanism underlying the downregulation of TCR surface expression by CD3ζ-ERR, we used intracellular staining of CD3 to evaluate the localization of TCR complexes within T cells. The results revealed a modest increase in intracellular CD3 MFI in CD3ζ-ERR T cells, showing an approximate 1.3-fold elevation that correlated positively with GFP expression levels (Figure 2A and B). Conversely, the surface expression of CD3 in CD3ζ-ERR T cells was significantly reduced compared to the control group (Figure 2C and D). These observations suggest that in CD3ζ-ERR T cells, TCR complexes that fail to reach the cell surface are likely sequestered within intracellular compartments, thereby supporting the intended functional design of the CD3ζ-ERR construct.

**Figure 2.**
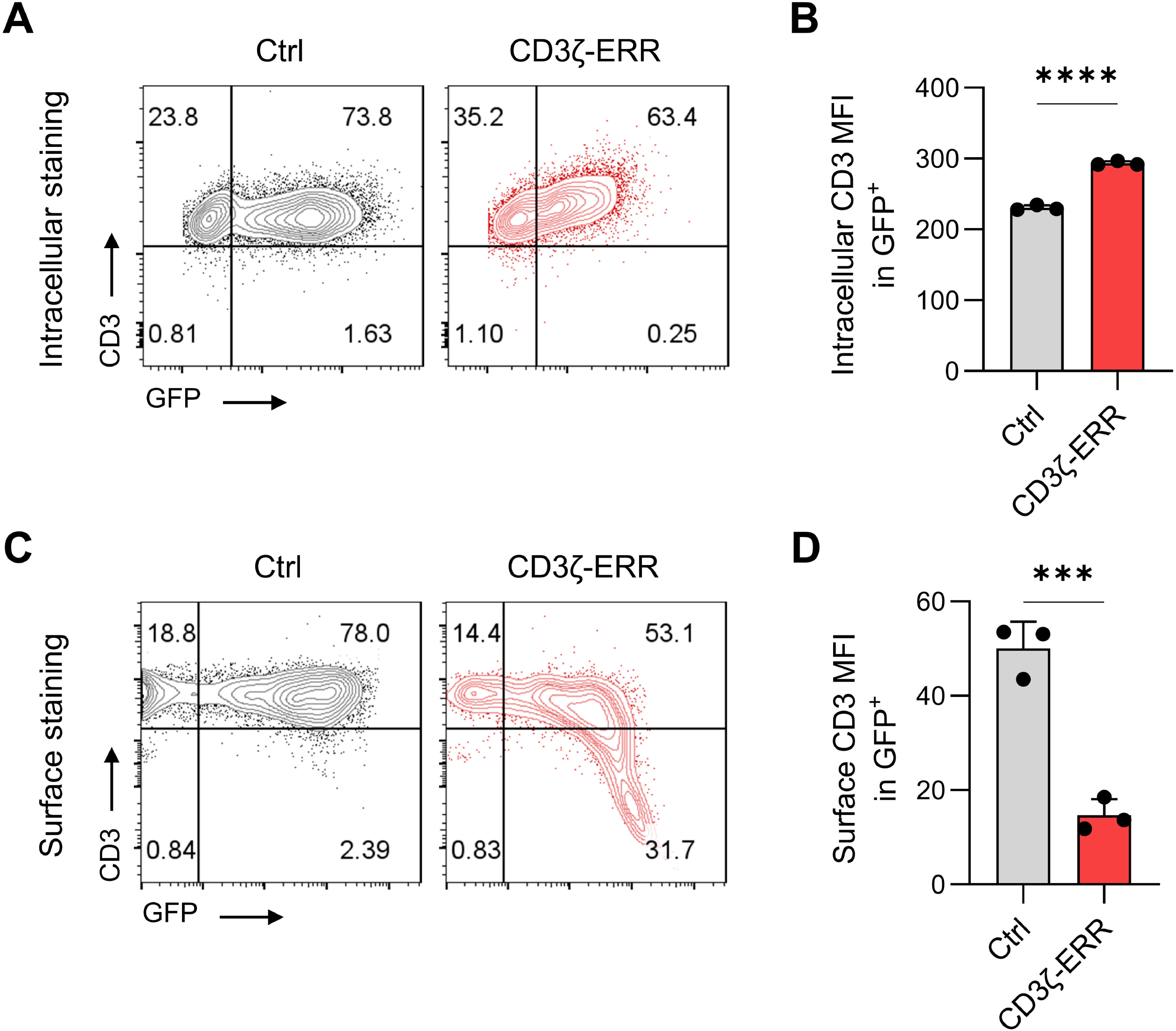
CD3ζ-ERR sequester CD3 intracellularly in T cells. **A**.Representative flow cytometry result of intracellular staining of CD3 showing slightly increasing CD3 staining intensity within correlation with GFP signal **B**. summary data for three biological replicates showing increased intracellular geometric mean fluorescence intensity (MFI) of intracellular CD3 staining in GFP+ CD3ζ-ERR T cells compared to control (Ctrl) GFP-FFluc-transduced T cells. **C**. Representative surface staining of CD3 showing reduced CD3 expression in CD3ζ-ERR T cells. **D**. The bar graph summarizing three biological replicates of CD3 surface staining on CD3ζ-ERR and Ctrl T cells showing reduced MFI of CD3 in GFP+ population The data are presented as mean ± SD, \*\*\*\**P* < 0.0001, *** *P* < 0.001, unpaired t test was used for statistical analysis.

### CD3ζ-ERR does not impair T cell proliferation, cell cycle dynamics, phenotype, or viability

To confirm that CD3ζ-ERR expression does not negatively impact T cell viability or proliferation, we examined the survival and proliferation of CD3ζ-ERR-transduced T cells during the ex-vivo expansion. We did not observe any significant difference between the GFP-FFluc control and CD3ζ-ERR T cell numbers during the expansion (Figure 3A). Also, CD8 staining on day 11 in transduced control and CD3ζ-ERR T cells revealed no significant difference suggesting that CD3ζ-ERR does not affect CD4:CD3 ratio (Figure 3B and C). Additionally, Cell cycle analysis on day 11 demonstrated no significant differences in the distribution of cells across the G1, S, G2, or subG1 phases (Figure 3D and E). Viability assessment using side and forward scatters, and PI staining also revealed no significant differences in viability of these cells (Figure 2F, G, and H). These findings indicate that CD3ζ-ERR T cells maintain comparable viability, proliferation, and cell cycle dynamics, with no significant shift in the CD8 ratio.

**Figure 3.**
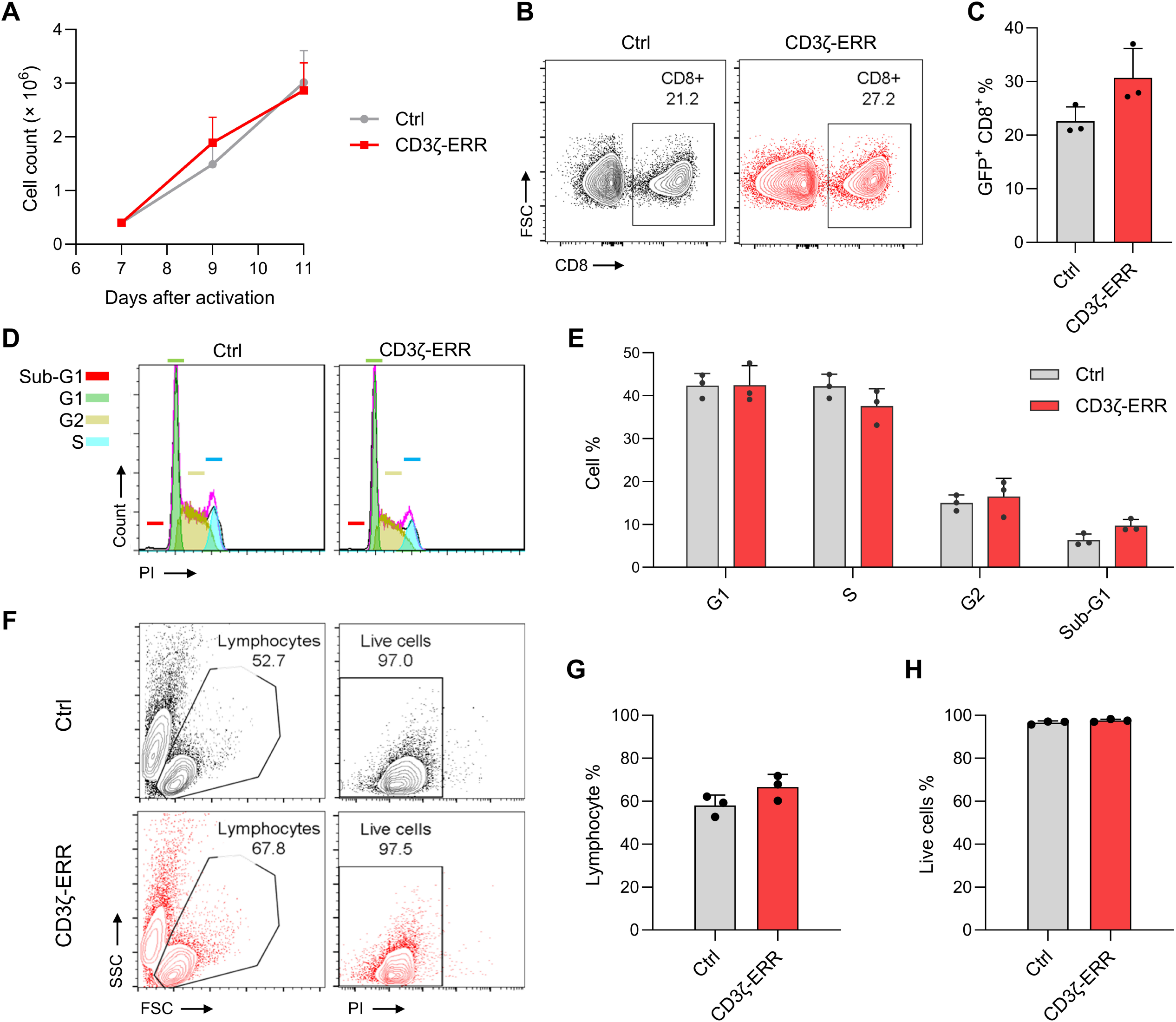
CD3ζ-ERR toxicity analysis on T cells. **A.**Line graph of cell counts. On day 9 and 11 cell counts were applied to reveal the amplification ratio. No difference was seen between the groups. **B**. Total cellular counts on day 11, which revealed no difference between groups. **C**. The scatter plot of CD8 expression. **D**. No difference in T cell CD8 phenotypes were seen between groups. **E**. Cell cycle curve analysis. **F**. The bar graph of different cell cycle stages on day 11 which shown no difference between groups. **G**. Scatter plot of first gating on FSC and SSC and second gating based on PI for live cell detection. **H**. The bar graph in lymphocyte gating which have no difference in groups. **I**. The bar graph in live cell gating which have no difference in groups. One-way ANOVA or unpaired t test applied and did not show any statically significant difference.

### CD3ζ-ERR T cells have deficient TCR-mediated activation

To confirm that suppression of CD3 expression on T cell surface by CD3ζ-ERR abolishes their ability to activate via the endogenous TCR, we assessed pan-T cell activation in the presence of agonistic anti-CD3 antibody (OKT3). We first sorted GFP^+^CD3^−^ T cells seven days post-transduction with CD3ζ-ERR, while control GFP-FFluc T cells were sorted based on GFP expression (Figure 4A, B, and C). The sorted T cells were exposed to plate-bound anti-CD3 for 48 hours, and T cell activation was evaluated by measuring CD69 expression, proliferation, and IL-2 secretion. The results indicated that CD3ζ-ERR T cells exhibited significantly lower CD69 expression than control T cells (Figure 4D and E). Additionally, cell counting indicated a reduced proliferation ratio in CD3ζ-ERR T cells (Figure 4F). IL-2 secretion levels measured in the media were also significantly lower in CD3ζ-ERR T cells (Figure 4G). These results confirm that the downregulation of TCR surface expression by CD3ζ-ERR leads to impaired TCR-mediated activity, even in the presence of a potent agonistic anti-CD3 antibody.

**Figure 4.**
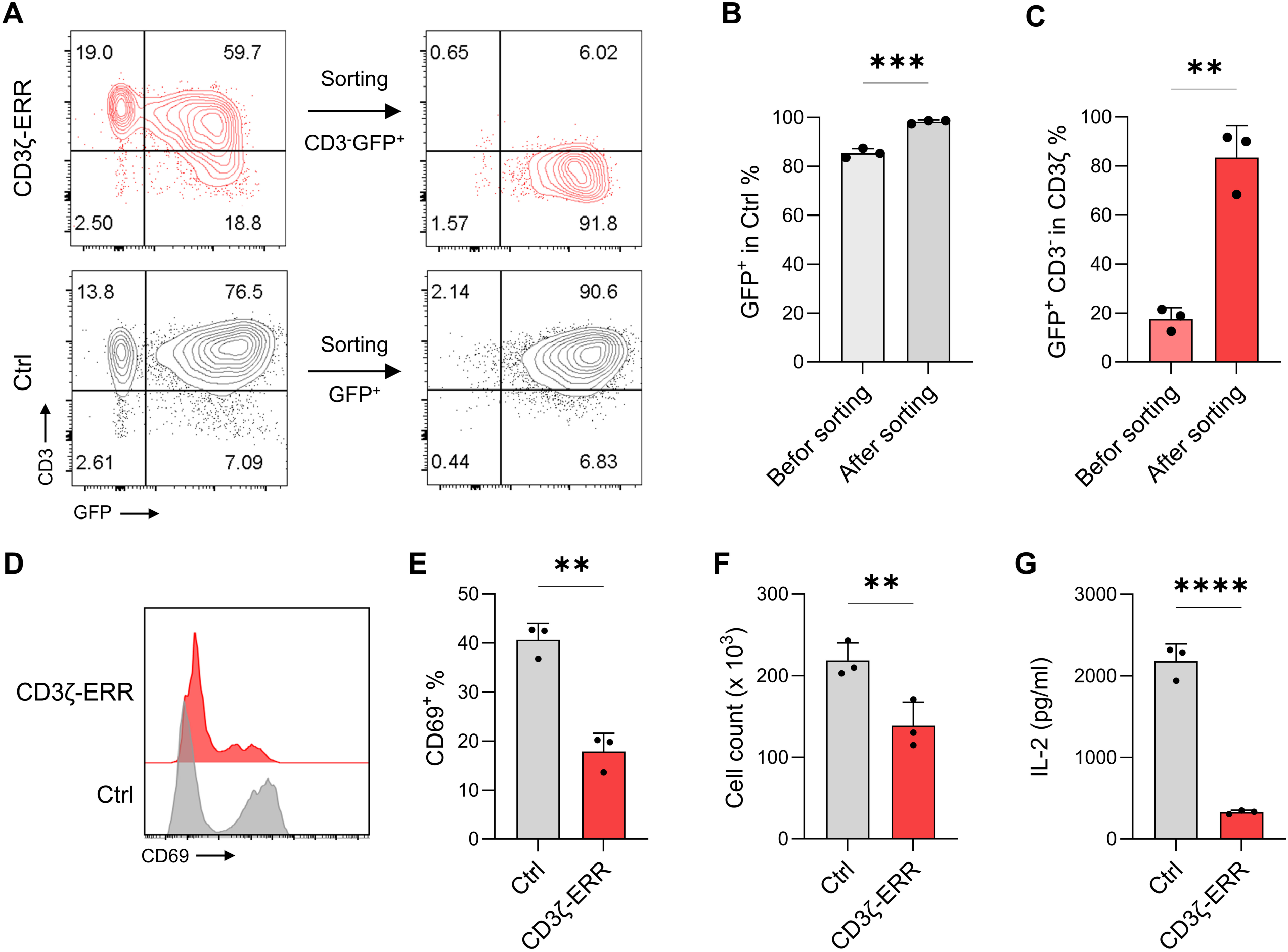
CD3ζ-ERR sorting, anti-CD3 activation. **A**. The scatter plot of CD3^+^ GFP^+^ expression before and after flow sorting. **B**. After sorting 98.27 % of Ctrl T cells are GFP^+^ **C**. After sorting 83.43 % of CD3ζ-ERR T cells are GFP^+^ CD3^-^ Which were significantly higher than before sorting **D**. The histogram graph of CD69 expression 48h after anti-CD3 activation. **E**. The CD69 expression revealed significant difference with Ctrl group due to downregulation of TCR in CD3ζ-ERR T cells **F**. The bar graph of cell counts which was significantly different between groups. **G**. IL-2 secretion significantly decreased in CD3ζ-ERR T cells after anti-CD3 activation. (The data are presented as mean ± SD, \*\*\*\**P* < 0.0001, *** *P* < 0.001, ** *P* < 0.01, * *P* < .05, unpaired *t*-test was applied).

### CD3ζ-ERR attenuates alloreactivity in ex vivo expanded T cells

To investigate the alloreactivity potential of the CD3^−^ sorted CD3ζ-ERR T cells, we conducted a co-culture experiment with allogenic PBMCs. We measured T cell proliferation and cytokine secretion 72 hours after starting the coculture to assess alloreactivity. The proliferation marker Ki67 was significantly reduced in CD3ζ-ERR T cells, showing a decrease of approximately 3.75-fold compared to control T cells (Figure 5A and B). Additionally, IFN-γ secretion was significantly lower in CD3ζ-ERR T cells (Figure 5C). These results showed that the impaired TCR-mediated activation in CD3^−^ CD3ζ-ERR T cells significantly diminished their alloreactive response.

**Figure 5.**
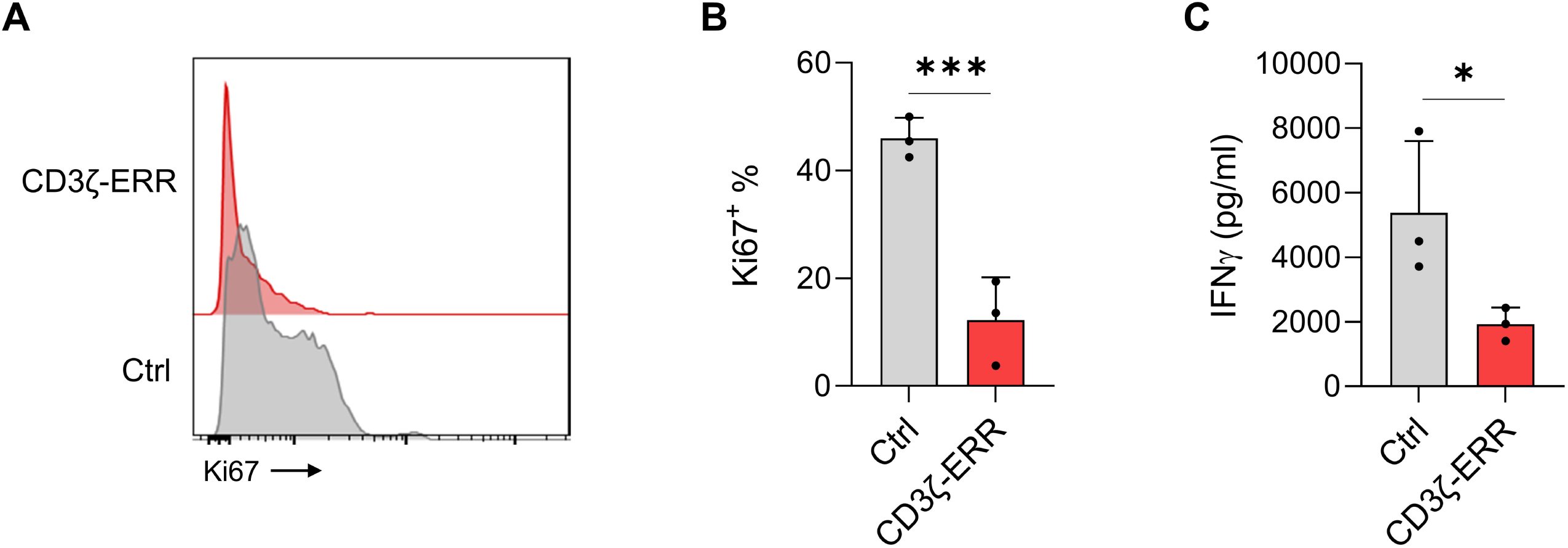
In vitro alloreactivity assay. **A**.The histogram of Ki67 72h after coculture with alloreactive PBMC. **B**. Ki67 was significantly higher in Ctrl group which related to proliferation response in alloreactivity condition. **C**. IFN-γ also was significantly high for Ctrl group that related to functional activity in alloreactivity assay. (The data are presented as mean ± SD, \*\*\*\**P* < 0.0001, *** *P* < 0.001, ** *P* < 0.01, * *P* < 0.05, unpaired *t*-test was applied).

### CD3ζ-ERR preserves the antitumor efficacy of CAR T cells

The application of CD3ζ-ERR in allogeneic CAR T cells necessitates verifying that its expression does not adversely affect CAR functionality. To assess this, we engineered CD19 CAR T cells to co-express CD3ζ-ERR and compared their antitumor activity with conventional CD19 CAR T cells. We first evaluated CAR-induced T cell activation by measuring CD69 expression following co-culture with CD19^+^ Raji cells for 24 hours. No significant differences in CD69 expression were observed between the CD3ζ-ERR CAR T cells and control CAR T cells, at both 1:1 and 20:1 effector to target (E:T) ratios (Figure 6A and B).

**Figure 6.**
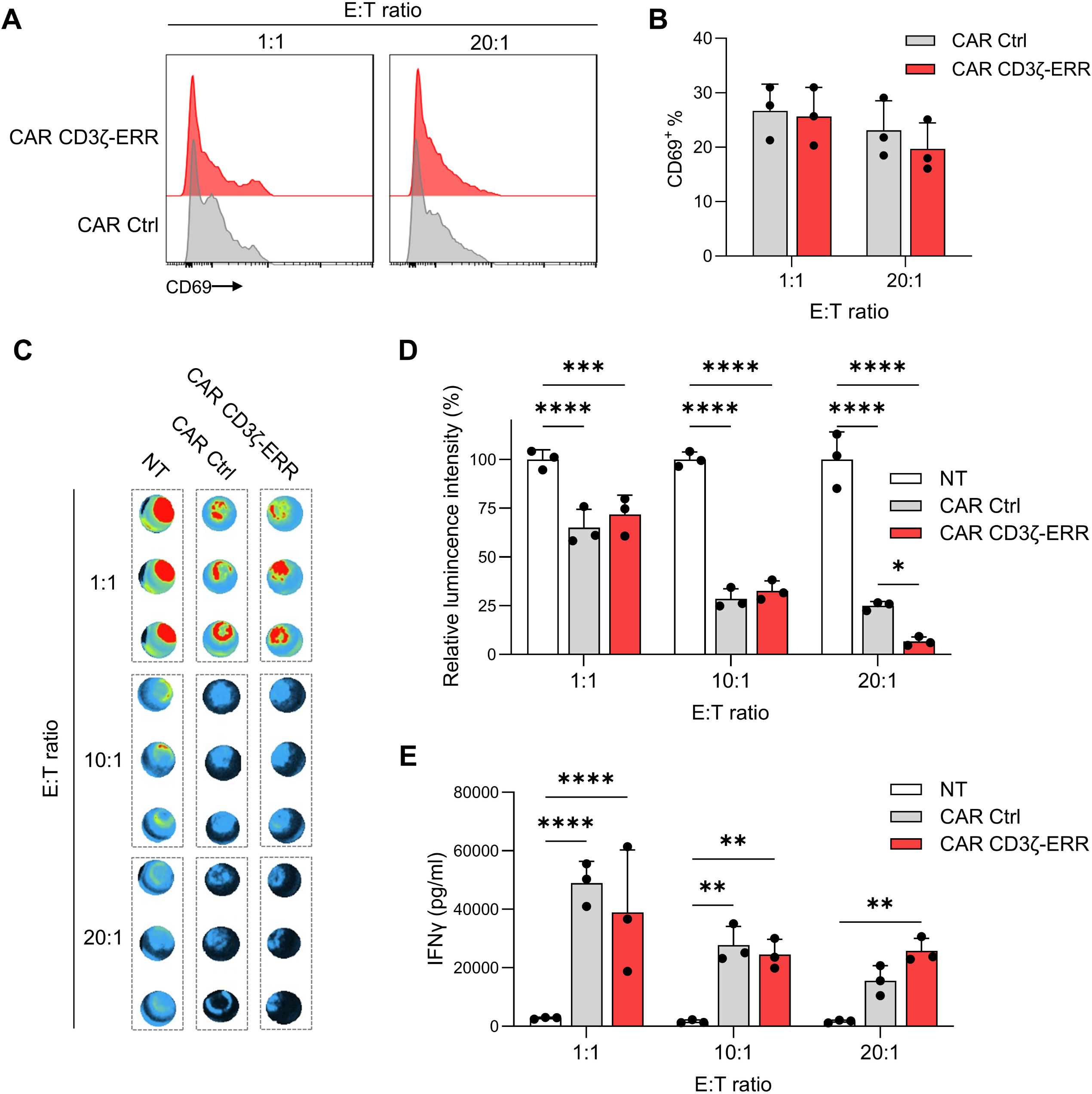
In vitro anti-tumor response of CAR CD3ζ-ERR T cells. **A**.The histogram of CD69 expression 24h after coculture with Raji cells. **B**. CD69 expression reveal no difference between CAR Ctrl and CAR CD3ζ-ERR T cells in E: T (1:1 and 20:1). **C, D**. Specific tumor cell lysis by CAR T cells. CAR Ctrl T cells, CAR CD3ζ-ERR T cells and untransduced T cells (NT) were co-cultured with ffluc-expressing Raji cells at different effector to target (E: T) ratios and after 24 hours of incubation cell lysis was measured in a luciferase-based assay. Both CAR groups significantly have higher lysis response compared to NT cells however, CAR groups have no significant difference with each other. **E**. IFN-γ also was significantly high in CAR groups in comparison with NT. (The data are presented as mean ± SD, \*\*\*\**P* < .0001, *** *P* < .001, ** *P* < .01, * *P* < .05, one-way ANOVA or unpaired t test were applied based on groups).

We then examined CAR-mediated cytotoxicity using a luciferase-based killing assay across E:T ratios of 1:1, 10:1, and 20:1. Both conventional CD19 CAR T cells and CD3ζ-ERR CD19 CAR T cells demonstrated significant tumor cell lysis at all tested ratios compared with untransduced T cells (Figure 6C and D). While cytotoxicity was similar between the two CAR T cell groups at 1:1 and 10:1 ratios, CD3ζ-ERR CAR T cells showed slightly enhanced tumor cell killing at the 20:1 ratio (Figure 6D). Additionally, we quantified IFN-γ secretion in these co-culture conditions, finding that both CAR T cell groups produced comparable levels of IFN-γ, which were significantly higher than the levels produced by untransduced T cells (Figure 6E). These findings indicate that CD3ζ-ERR does not interfere with CAR-mediated antitumor responses, supporting its potential use in developing potent allogeneic CAR T cell therapies.

## Discussion

In this study, we developed an efficient and straightforward method for the downregulation of endogenous TCR expression in T cells, without the need for genome editing. By introducing the CD3ζ-ERR construct via viral transduction, we successfully disrupted the membrane trafficking of the TCR complex, thereby preventing its surface expression on T cells. We demonstrated that CD3ζ-ERR effectively reduces endogenous TCR-mediated activation and alloreactivity, while maintaining the robust CAR-mediated tumor-killing function when co-expressed in T cells.

Allogeneic αβ CAR T cells engineered through gene editing technologies have shown substantial promise in both preclinical and clinical contexts.^18^ However, these approaches necessitate extensive quality control to avoid off-target effects, rendering the manufacturing process complex and challenging.^19^ Furthermore, recent studies have raised significant concerns regarding genome integrity, particularly due to on-target activities of genome editing that may result in persistent chromosome loss and large unintended deletions in human T cells and CAR T cells designed for clinical use.^10, 12, 20^ While genome editing remains a feasible and promising strategy for the development of allogeneic CAR T cells, our non-genome-editing approach eliminates these safety risks and simplifies the manufacturing process. Additionally, our method does not require substantial modifications to current viral vector-based manufacturing protocols, making it more compatible with existing clinical-grade T cell production systems. The relatively short genetic sequence of the CD3-ERR construct (200 bp-670 bp) allows it to be easily combined with the CAR gene within a single retroviral vector, facilitating integration into existing GMP-grade large-scale manufacturing processes with minimal adjustments.

To design the CD3-ERR constructs, we incorporated an ERR signal, previously shown to effectively sequester chimeric proteins within the ER,^14, 21^ into truncated or full-length CD3ζ and CD3ε subunits of the TCR complex. Given that these subunits each contribute two copies to the TCR complex, the higher stoichiometry increases the likelihood of transgenic incorporation into endogenous TCR assemblies. Not surprisingly, we observed an inverse correlation between transgene expression (GFP intensity) and CD3 surface expression across all CD3-ERR constructs, indicating a competitive interaction between the CD3-ERR constructs and their endogenous counterparts. Notably, the CD3ζ-ERR construct, which exhibited the highest transgene expression, also achieved the greatest overall TCR suppression, comparable to the efficiency reported for protein expression blockers (PEBLs) that use anti-CD3ε scFv to prevent TCR complex surface trafficking.^14^

Our results establish proof of concept for the efficacy of CD3ζ-ERR in effectively diminishing endogenous TCR-mediated alloreactivity without impairing CAR-mediated antitumor activity. However, further investigations are necessary to fully assess the safety profile of CD3ζ-ERR CAR T cells before their transition into clinical applications. Although CD3ζ-ERR expression resulted in near-complete TCR suppression in a significant portion of the transduced T cell population, some cells continued to express varying levels of TCR, necessitating additional cell sorting steps. Given the competition-based mechanism of action, combining CD3ζ-ERR with other TCR downregulation strategies, such as RNA interference, which introduces minimal genetic payload,^13, 18^ could enhance the efficiency of TCR suppression and warrants further exploration.

In addition to strategies that target TCR disruption, HLA-matching has also been investigated as a method to mitigate GvHD in allogeneic CAR T cell therapy. Clinical studies have suggested a lower-than-expected risk of GvHD when CD28-based CAR T cells are infused into HLA-matched patients.^22-25^ However, GvHD remains a significant concern, with reports of grade II GvHD in two of three patients treated with allogeneic CAR T cells, and grade II/III GvHD in three of six patients who received haploidentical CAR T cells.^26, 27^ These findings suggest that combining haploidentical approaches with TCR downregulation could be effective in reducing alloreactivity and enhancing the safety of allogeneic CAR T cell therapies.

In summary, our study demonstrates the potential of CD3ζ-ERR as a novel strategy for downregulating TCR expression in T cells without the need for genome editing. This approach effectively preserves CAR-mediated antitumor activity while mitigating the risk of alloreactivity, offering a promising alternative for the scalable production of safe and effective allogeneic CAR T cells. Continued investigation is warranted to enhance the robustness and clinical readiness of this strategy.

## Acknowledgement

We thank all staff and technicians at the Royan Institute for Stem Cell Biology and Technology for their technical assistance. This study was supported by a grant from Royan Institute (Grant No. IR.ACECR.ROYAN.REC.1401.116).

## Data Availability Statement

The data generated during this study is available upon reasonable request to the corresponding authors.

